# SARS-CoV-2 wildlife surveillance in Ontario and Québec, Canada

**DOI:** 10.1101/2021.12.02.470924

**Authors:** Janet E. Greenhorn, Jonathon D. Kotwa, Jeff Bowman, Laura Bruce, Tore Buchanan, Peter A. Buck, Antonia Dibernardo, Logan Flockhart, Marianne Gagnier, Aaron Hou, Claire M. Jardine, Stephane Lair, L. Robbin Lindsay, Ariane Masse, Pia K. Muchaal, Larissa A. Nituch, Angelo Sotto, Brian Stevens, Lily Yip, Samira Mubareka

## Abstract

**Background:** Severe acute respiratory syndrome coronavirus 2 (SARS-CoV-2), the virus responsible for the COVID-19 pandemic, is capable of infecting a variety of wildlife species. Wildlife living in close contact with humans are at an increased risk of SARS-CoV-2 exposure and if infected have the potential to become a reservoir for the pathogen, making control and management more difficult.

**Objective:** To conduct SARS-CoV-2 surveillance in urban wildlife from Ontario and Québec, Canada, increasing our knowledge of the epidemiology of the virus and our chances of detecting spillover from humans into wildlife.

**Methods:** Using a One Health approach, we leveraged activities of existing research, surveillance, and rehabilitation programs among multiple agencies to collect samples from 776 animals from 17 different wildlife species between June 2020 and May 2021. Samples from all animals were tested for the presence of SARS-CoV-2 viral RNA, and a subset of samples from 219 animals across 3 species (raccoons, *Procyon lotor*; striped skunks, *Mephitis mephitis*; and mink, *Neovison vison*) were also tested for the presence of neutralizing antibodies.

**Results:** No evidence of SARS-CoV-2 viral RNA or neutralizing antibodies was detected in any of the tested samples.

**Conclusion:** Although we were unable to identify positive SARS-CoV-2 cases in wildlife, continued research and surveillance activities are critical to better understand the rapidly changing landscape of susceptible animal species. Collaboration between academic, public and animal health sectors should include experts from relevant fields to build coordinated surveillance and response capacity.

## Introduction

The severe acute respiratory syndrome coronavirus 2 (SARS-CoV-2) is responsible for the global COVID-19 pandemic and has been maintained through human-to-human transmission. However, humans are not the only species susceptible to infection. Over the course of the current pandemic, a range of domestic and wild animal species have been reported to either be naturally infected with SARS-CoV-2 or susceptible to the virus in experimental infections (1, 2, 3). Others have been identified as potential hosts based on sequence analysis of the host cell receptor of SARS-CoV-2, angiotensin 1 converting enzyme 2 (ACE2), and predicted binding affinity (4, 5).

Many wild animal species thrive in the ecological overlap with humans and are thus at an increased risk of being exposed to SARS-CoV-2 (6). Several of these peri-domestic species have been experimentally shown to become infected with and shed SARS-CoV-2 (7, 8). SARS-CoV-2 infection has also been reported in wild or free-ranging animals that have been naturally exposed, including American mink (*Neovison vison*; 9) and, more recently, white-tailed deer (*Odocoileus virginianus*; 10, 11).

The concept of One Health recognizes that human health and animal health are interdependent (12). The spillover of virus from humans or domestic animals into wildlife is concerning not only due to the possible deleterious effects on wildlife, but because these wild populations have the potential to act as reservoirs for SARS-CoV-2. Diseases that have an animal reservoir are inherently much more difficult to control and the spread of SARS-CoV-2 through animal populations could further contribute to the development of variants of concern (VoCs), potentially undermining the efficacy of medical countermeasures such as antivirals and vaccines (13, 14). Additionally, people who have close contact with wildlife, such as biologists, wildlife rehabilitators, and hunters and trappers, may be at higher risk of being exposed to the virus and of facilitating its spread among wildlife. The impact of SARS-CoV-2 infection on wildlife health is not fully understood. Early detection of any spillover is therefore critical to preventing and addressing these concerns.

Given the risk of reverse-zoonotic SARS-CoV-2 transmission and our lack of knowledge of the virus in local wildlife, there was an urgent need to elucidate the epidemiology of the virus at the human-wildlife interface to help wildlife management and public health officials better communicate risk and plan management strategies. We therefore conducted SARS-CoV-2 surveillance in wildlife across Ontario and Québec, Canada, with a major focus on the southern regions of both provinces. These areas have high human population densities and include major urban centres such as Toronto and Montréal. Incidences of COVID-19 peaked in Montréal and the surrounding regions in early January 2021, with rates exceeding 400 cases per 100,000 population in Montréal and Laval (15). Incidences in Toronto and the surrounding regions peaked in April 2021, with case rates in the City of Toronto and Peel also exceeding 400 per 100,000 population (15).

## Methods

Many experts have recommended a One Health approach for animal SARS-CoV-2 testing, which balances concerns for both human and animal health and is based on knowledge of experts in both fields (16, 17). As such, our work was conducted through consultation and cooperation among a wide variety of agencies: the Public Health Agency of Canada (PHAC), the Ontario Ministry of Northern Development, Mines, Natural Resources and Forestry (NDMNRF), le Ministère des Forêts, de la Faune et des Parcs du Québec (MFFP), the Canadian Wildlife Health Cooperative (CWHC), the Ontario Ministry of Agriculture, Food, and Rural Affairs (OMAFRA), the Canadian Food Inspection Agency (CFIA), the Western College of Veterinary Medicine, the Granby Zoo, the National Microbiology Laboratory (NML) of PHAC, and Sunnybrook Research Institute (SRI). We focussed our surveillance primarily on animals from urban areas or those with a case history of close contact with people since these animals would be at the highest risk of exposure to people infected with SARS-CoV-2. All samples for testing were collected between June 2020 and May 2021 through pre-existing partnerships or over the course of other research, surveillance, or rehabilitation work (Table 1).

**Table 1:**
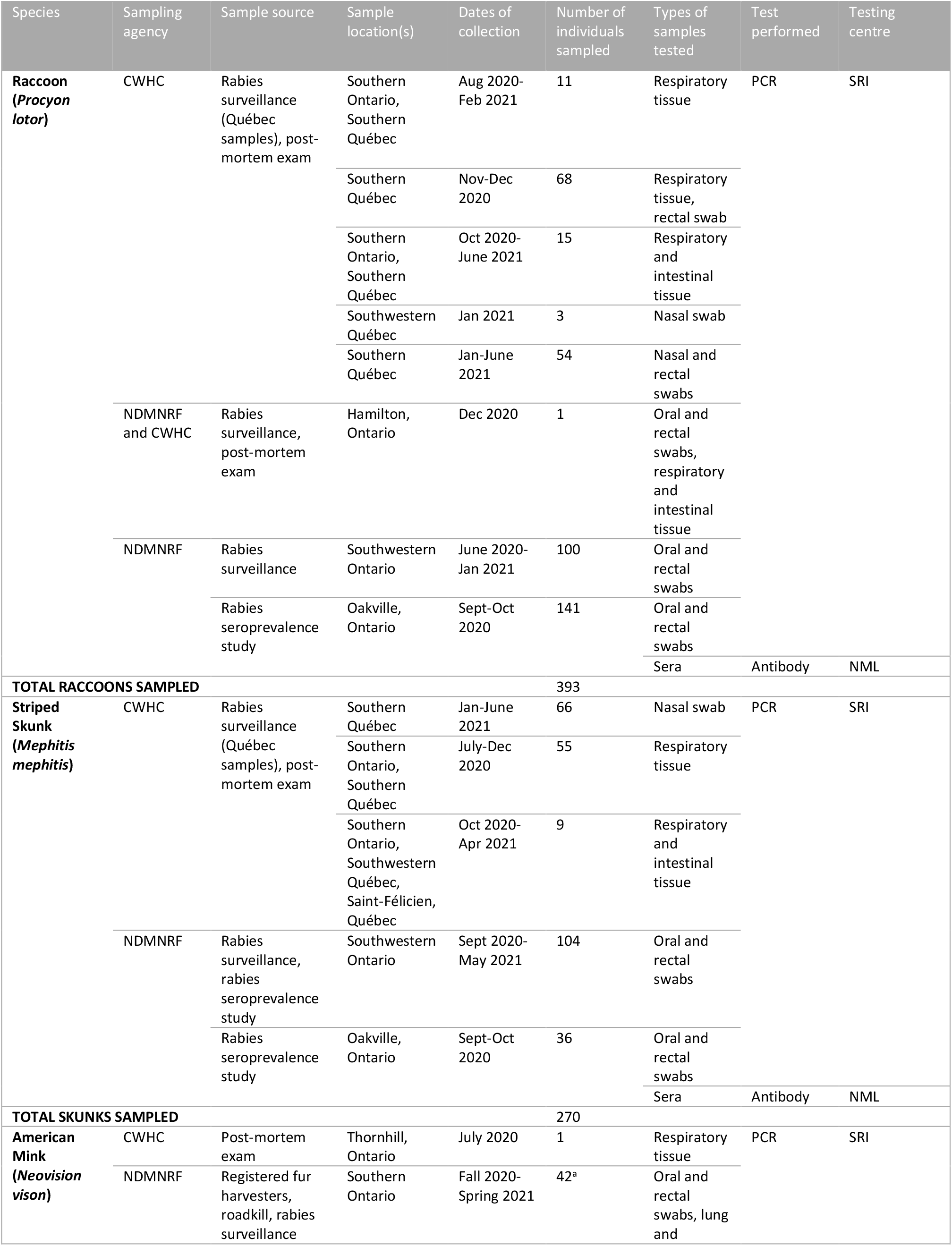
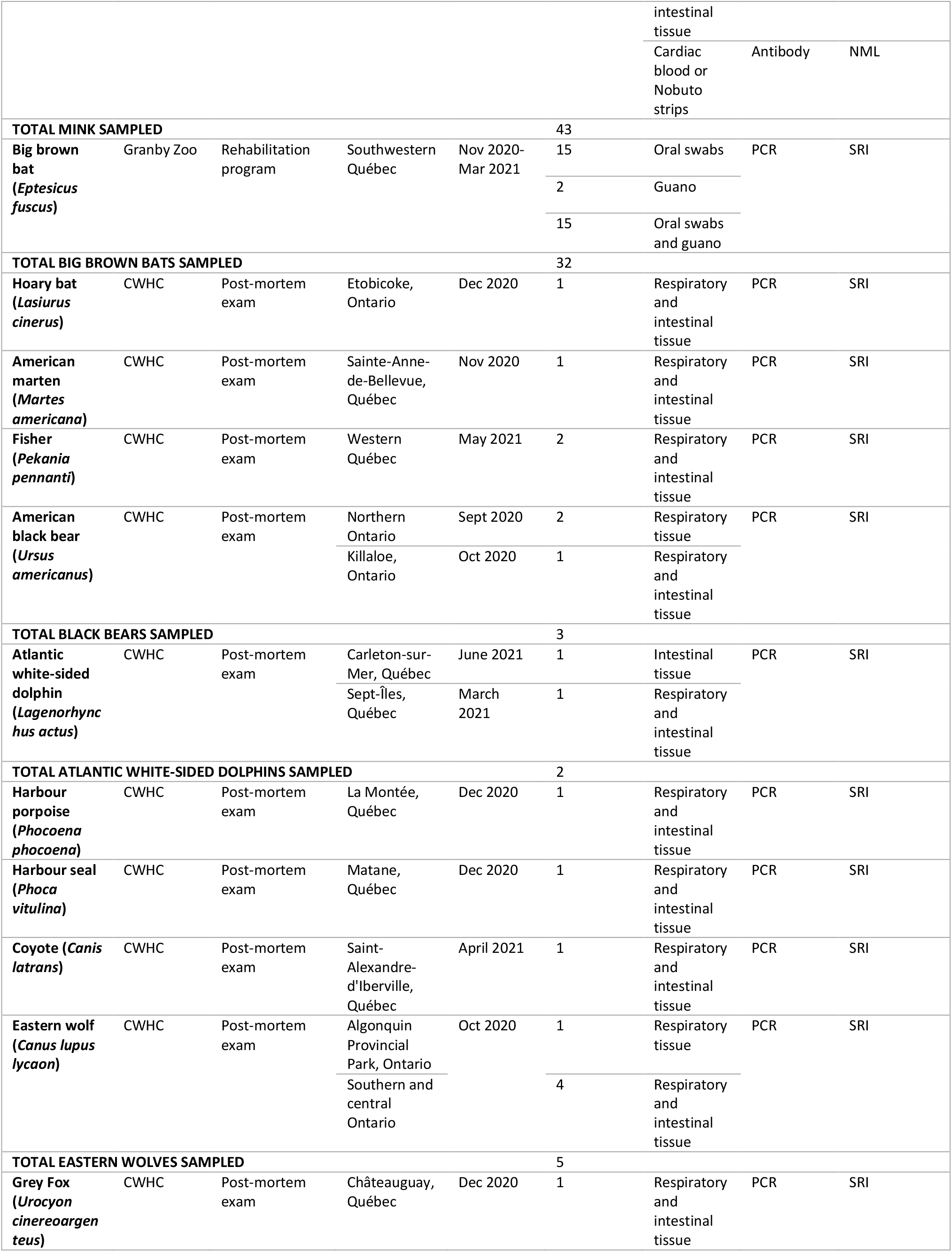
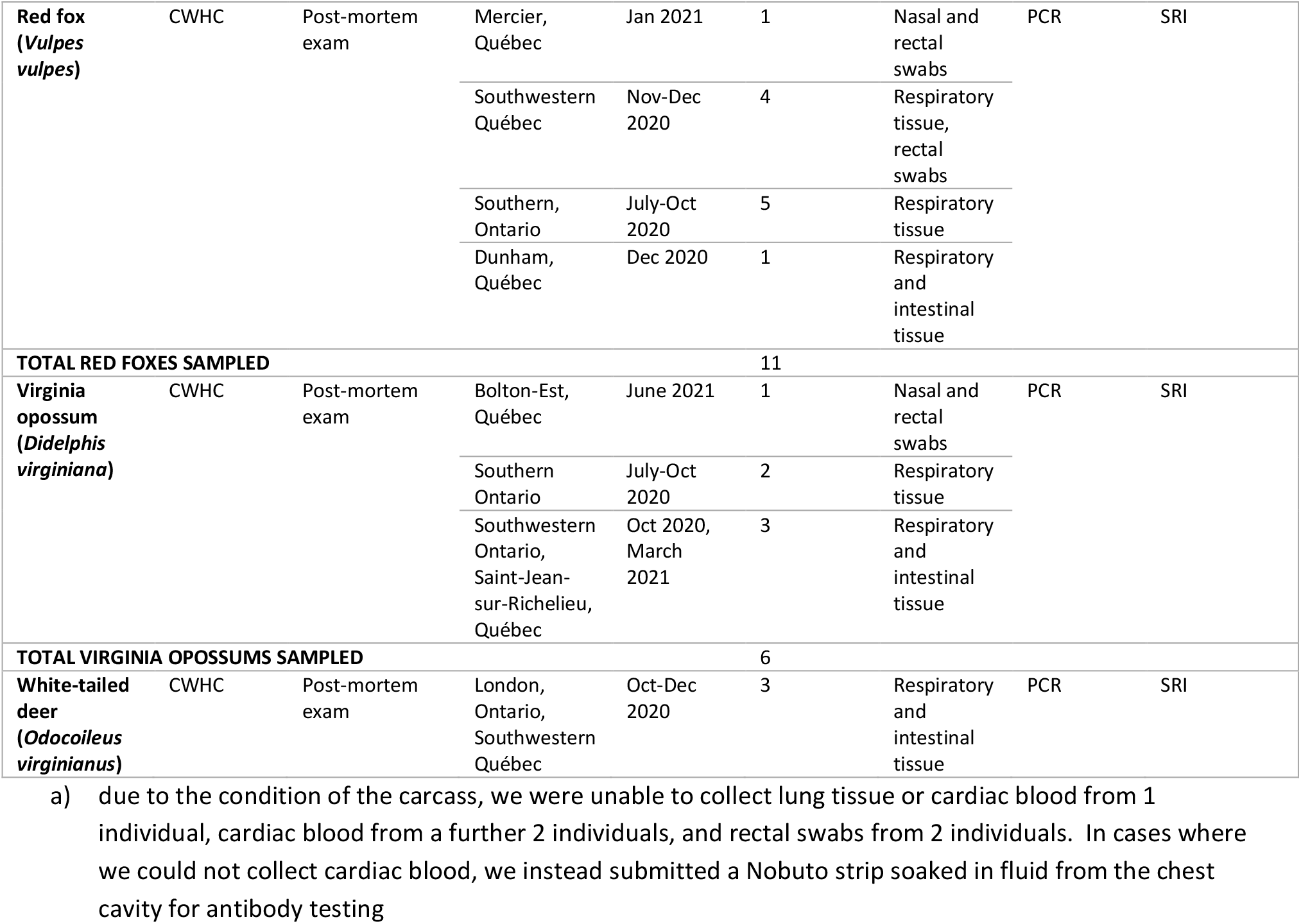
Metadata for 776 animals from Ontario and Québec screened for SARS-CoV-2.

### Raccoons and skunks

Raccoons (*Procyon lotor*) and striped skunks (*Mephitis mephitis*) are peri-domestic species that are good candidates for reverse-zoonotic disease surveillance due to their high density in urban areas and their frequent close contact with people, pets, and refuse. They are also subject to ongoing rabies surveillance operations in both Ontario and Québec, making them easy to sample. In Ontario, wildlife rabies surveillance and testing are conducted by the NDMNRF on roadkill, animals found dead for other reasons, and deceased sick or strangely acting wildlife. Submissions are received mainly from southwestern Ontario, and most animals received by the program and subsequently sampled and tested for SARS-CoV-2 came from urban centres within this region (Figure 1). In Québec, a similar wildlife rabies surveillance program is coordinated by the MFFP and testing and other post-mortem examinations are performed by the Québec CWHC. As was the case in Ontario, animals sampled by the Québec CWHC for SARS-CoV-2 testing came mainly from urban areas (Figure 1). The Ontario CWHC laboratory also contributed a small number of raccoon and skunk samples from animals submitted to them for post-mortem examination. Carcasses were sampled using a combination of oral, nasal, and rectal swabs, respiratory tissue, and intestinal tissue (Table 1). Swabs were stored in individual 2 mL tubes with ∼1 mL of universal transport medium (UTM; Sunnybrook Research Institute) and 30-60 mg tissue samples were stored dry in tubes.

**Figure 1:**
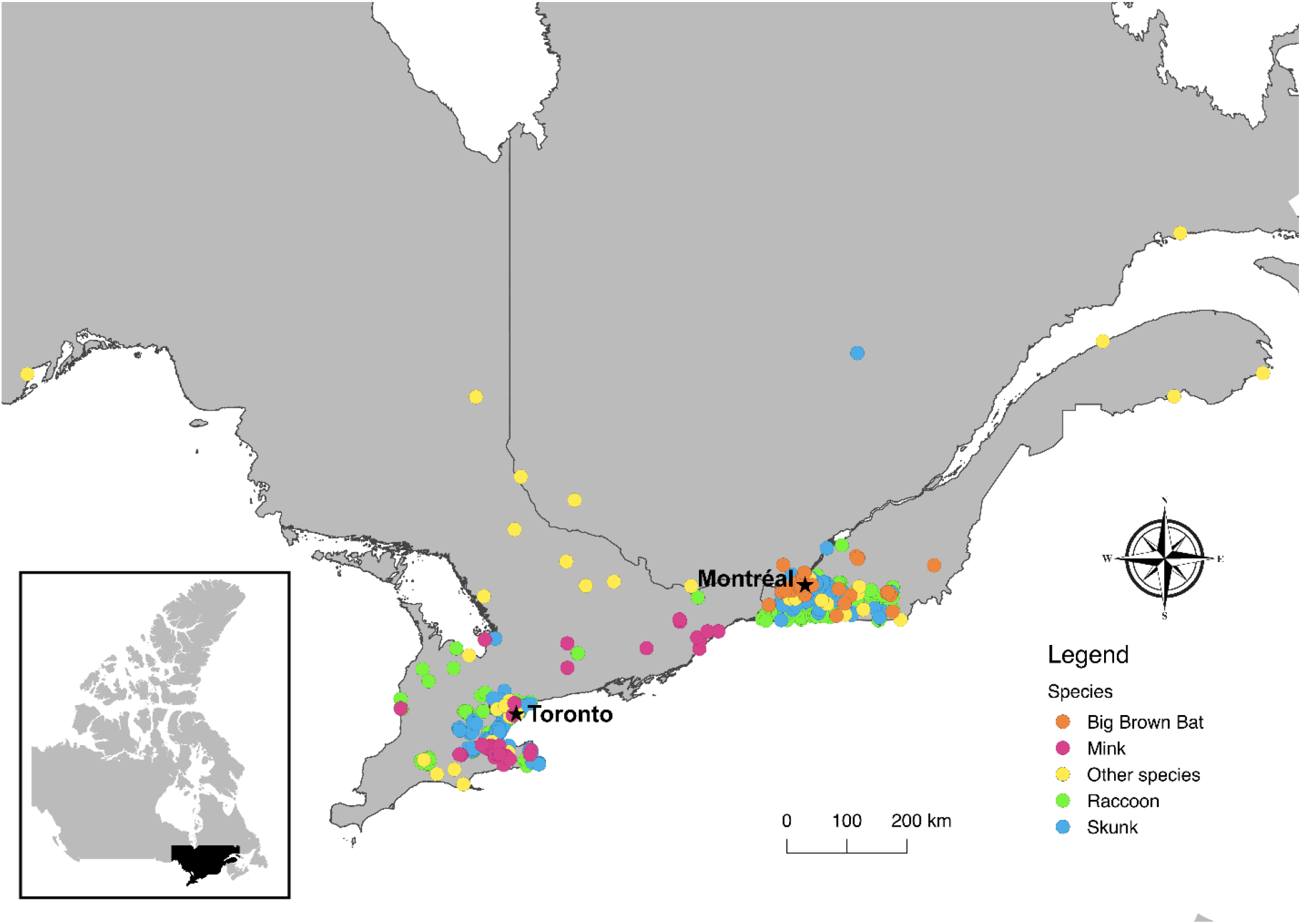
Original locations of animals submitted for SARS-CoV-2 testing (N=776)

Additionally, samples were collected from live raccoons and skunks during an annual seroprevalence study conducted by the NDMNRF in Oakville, Ontario to assess the effectiveness of rabies vaccine baiting (NDMNRF Wildlife Animal Care Committee Protocol #358). Animals were captured in live traps and transported to a central processing station where they were anaesthetized. Oral and rectal swabs were collected for PCR testing. Blood was drawn from the brachiocephalic vein and 0.2-1.0 mL of sera was collected for antibody testing. Following reversal and successful recovery, animals were returned to their point of capture and released.

### Mink

Instances of SARS-CoV-2 infection in mink have already been identified in multiple countries, including Canada, and infected farmed mink have proven capable of passing the virus to naïve conspecifics, humans, and domestic and feral companion animals (18, 19, 20, 21, 22). At the time of writing no mink farm outbreaks have been reported in Ontario or Québec, but mink farms in Ontario have previously been shown to act as points of infection for other viruses (e.g. Aleutian Mink Disease), which can spread to wild mink populations (23).

The majority of mink carcasses we sampled for SARS-CoV-2 were submitted to the NDMNRF by licensed fur harvesters through a collaboration with the Ontario Fur Managers Federation. The NDMNRF staff collected oral and rectal swabs, lung tissue, and intestinal tissue from the carcasses, as well as cardiac blood samples via cardiac puncture for antibody testing. If blood could not be obtained from the heart, fluid was collected from the chest cavity on a Nobuto filter strip (Advantec MFS, Inc, Dublin, CA, USA). Nobuto strips were allowed to air dry, then placed in individual coin envelopes.

### Big brown bats

Bats are known carriers of coronaviruses (24, 25, 26). As such, concerns have been raised over the possible susceptibility of North American bats to SARS-CoV-2 (27). Species such as the big brown bat (*Eptesicus fuscus*) frequently roost in buildings, which brings them into close contact with people and increases the likelihood of SARS-CoV-2 exposure. Big brown bat oral swabs and guano samples for SARS-CoV-2 PCR testing were collected by staff at the Granby Zoo, which runs a rehabilitation program over the winter to care for bats that have been disturbed during their hibernation. Guano samples were stored dry in 2 mL tubes.

### Other species

Other samples for SARS-CoV-2 PCR testing were obtained opportunistically through the Ontario and Québec regional CWHC laboratories, which receive a wide variety of wildlife species for post-mortem examination (Table 1). Animals were selected for sampling based on potential for SARS-CoV-2 infection. This could be due to urban habitat, human contact, or to predicted species susceptibility based on prior research. The number and type of samples collected varied by carcass and depended on carcass condition (Table 1).

### RNA Extraction

RNA extraction and PCR testing were performed at the SRI in Toronto, Ontario. All swab, tissue, and guano samples were stored at -80 °C prior to testing. For oral, rectal, or nasal swab samples, RNA extractions were performed using 140 µL of sample via the QIAmp viral RNA mini kit (Qiagen, Mississauga, ON, Canada) or the Nuclisens EasyMag using Generic Protocol 2.0.1 (bioMérieux Canada Inc., St-Laurent, QC, Canada) according to manufacturer’s instructions; RNA was eluted in 50 µL. RNA from 80 mg of guano samples were extracted via the QIAmp viral RNA mini kit and eluted in 40 µL. Tissue samples were thawed, weighed, minced with a scalpel, and homogenized in 600 µL of lysis buffer using the Next Advance Bullet Blender (Next Advance, Troy, NY, USA) and a 5 mm stainless steel bead at 5 m/s for 3 minutes. RNA from 30 mg tissue samples was extracted via the the RNeasy Plus Mini kit (Qiagen, Mississauga, ON, Canada) or the Nuclisens EasyMag using Specific Protocol B 2.0.1; RNA was eluted in 50 µL. All extractions were performed with a negative control.

### SARS-CoV-2 PCR analysis

Reverse-transcription polymerase chain reaction (RT-PCR) was performed using the Luna Universal Probe One-Step RT-qPCR kit (NEB). Two gene targets were used for SARS-CoV-2 RNA detection: the 5’ untranslated region (UTR) and the envelope (E) gene. The cycling conditions were: 1 cycle of denaturation at 60 °C for 10 minutes then 95 °C for 2 minutes followed by 44 amplification cycles of 95°C for 10 seconds and 60°C for 15 seconds. Quantstudio 3 software (Thermo Fisher Scientific Inc., Waltham, MA, USA) was used to determine cycle thresholds (Ct). All samples were run in duplicate and samples with Cts <40 for both gene targets in at least one replicate were considered positive.

### Antibody testing

Antibody testing was performed on cardiac blood, chest cavity fluid and serum samples at the NML in Winnipeg, Manitoba. All samples were stored at -20 °C prior to testing. Cardiac blood samples were collected onto Nobuto filter strips (Advantec MFS, Inc, Dublin, CA, USA; Fisher Scientific, Waltham, MA, USA) by saturating the length of the strip with 100 µl of blood. To obtain the 1:9 dilution required for testing, saturated Nobuto strips were cut into 4-5 pieces and placed into a 2 mL tube containing 360 µl phosphate buffered saline (PBS) pH 7.4 containing 0.05% Tween 20 and eluted overnight at 4 ⁰C. Nobuto strips collected from chest cavity fluid were processed in the same way, whereas serum samples were diluted 1:9 with Sample Dilution Buffer. Samples were mixed by vortexing and tested using the GenScript cPass™ SARS-CoV-2 Neutralization Antibody Detection Kit (GenScript USA, Inc. Piscataway, NJ, USA) according to the manufacturer’s protocol.

Briefly, 60 µl of a sample was added to 60 µl HRP-conjugated RBD solution and incubated at 37 ⁰C for 30 minutes. A 100 µl aliquot of the mixture was transferred to the ELISA microwell test plate and incubated at 37 ⁰C for 15 minutes. Microwells were washed 4 times with 260 µl wash buffer then 100 µl TMB substrate was added to each well. Following a 20 minute incubation in the dark at room temperature, 50 µl of Stop Solution was added to each well. Absorbance was read immediately at 450 nm.

Each assay plate included positive and negative controls that met required quality control parameters. Percentage inhibition was calculated for each sample using the following equation:

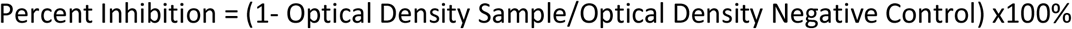

Samples with greater than or equal to 30% inhibition were considered positive for SARS-CoV-2 neutralizing antibodies.

## Results

We tested 776 individual animals from 17 different wildlife species for SARS-CoV-2. These animals were collected primarily from urban areas in southern Ontario and Québec between June 2020 and May 2021 (Table 1). We found no evidence of SARS-CoV-2 viral RNA in any of the tested samples and no evidence of neutralizing antibodies in a subset of 219 individuals (141 raccoons, 36 striped skunks, 42 mink).

## Discussion

Our study did not detect any spillover of SARS-CoV-2 to wildlife in Ontario and Québec. Raccoons and skunks were the most commonly tested species. Results from experimental studies have suggested these species may be susceptible to SARS-CoV-2, but the lack of and low quantity of infectious virus from raccoons and skunks, respectively, suggest they are an unlikely reservoir for SARS-CoV-2 in the absence of viral adaptations (7, 8). Similarly, a recent challenge study with big brown bats found that they are resistant to SARS-CoV-2 infection and do not shed infectious virus (28). Conversely, mink are susceptible to SARS-CoV-2 infection, but we did not detect evidence of SARS-CoV-2 in any of the mink sampled. While this could be attributed to our low effective sample size, to date SARS-CoV-2 has been infrequently detected in wild mink populations globally. It should be noted, however, that the abovementioned experimental studies on raccoons, skunks, and big brown bats were conducted using parental SARS-CoV-2. The susceptibility of these species to VoCs is presently not known and may differ from susceptibility to the parental strain (29). Additionally, challenge studies assessing susceptibility tend to be conducted on small numbers of young, healthy individuals, so results may not be reflective of the full range of possible responses to infection in the wild.

As the pandemic progresses, new evidence is emerging on susceptible wildlife that may act as competent reservoirs for the virus. For example, white-tailed deer are now considered a highly relevant species for SARS-CoV-2 surveillance in light of their experimentally determined susceptibility as well as evidence of widespread exposure to the virus via antibody and PCR testing across the northeastern USA (10, 11, 30). Continued surveillance efforts should be adaptive and include targeted testing of highly relevant species as they are identified. In Ontario and Québec, these would include mink, white-tailed deer, and deer mice (*Peromyscus maniculatus*; 7, 31). Continuing to include less susceptible species remains important given ongoing viral genomic plasticity and changing host range of VoCs.

## Limitations

There are several limitations for this study that need to be acknowledged. First, the majority of our SARS-CoV-2 testing was done by RT-PCR, which is only capable of detecting active infection. Antibody testing, which identifies resolved infection or exposure, is more likely to find evidence of SARS-CoV-2 in surveillance studies since results are less dependent on timing of sample collection. Antibody testing typically requires samples from live animals or fresh carcasses, which limited our ability to use it. However, the testing performed allowed for test validation in raccoons, skunks, and mink which may facilitate more antibody testing in future. Second, the type of samples we collected may also have limited our ability to detect SARS-CoV-2 infection. Viral replication can vary among tissue types and therefore some tissues are more optimal for viral RNA detection than others (1). In the present work, animals were sampled opportunistically as a part of pre-existing surveillance efforts, research, and rehabilitation programs and we were not able to consistently collect the same sample sets from all animals. Additionally, the sample types were from live animals and carcasses and not optimized; certain sample types were sometimes unavailable (e.g. tissue samples from live animals) or were not sufficient for collection.

## Conclusion

A One Health approach is critical to understanding and managing the risks of an emerging zoonotic pathogen such as SARS-CoV-2. We leveraged activities of existing surveillance, research, and rehabilitation programs and expertise from multiple fields to efficiently collect and test 1,690 individual wildlife samples. The absence of SARS-CoV-2-positive wildlife samples does not exclude spillover from humans to Canadian wildlife, given the limitations cited above. Continued research in this area is both important and pressing, particularly as novel VoCs emerge. Public and animal health sectors should continue to work collaboratively with academic and government partners to help prevent the spread of SARS-CoV-2 from people to wildlife, monitor for spillover, and address any issues should they arise. There is an urgent need for a coordinated wildlife surveillance program for SARS-CoV-2 in Canada. This approach will help protect the health of both Canadians and wildlife, now and in the future.

## Author’s Statement

JEG, JDK, JB, TB, PAB, LF, MG, CMJ, AM, PKM, LAN, SM - conceptualization

JEG, LB, MG, CMJ, SL, AM, BS - sample collection and coordination

JDK, AD, AH, LRL, AS, LY, SM – sample testing

JEG, JDK - resources

JEG, JDK, AD, LF – writing, original draft

JEG, JDK, JB, LB, TB, PAB, AD, LF, MG, AH, CMJ, SL, LRL, AM, PKM, LAN, AS, BS, LY, SM - writing, review and editing

JB, TB, PAB, PKM – funding acquisition

JEG and JDK contributed equally to this work.

## Competing Interests

None.

## Acknowledgements

The authors wish to thank B. Pickering and J. Tataryn for facilitating the inter-agency partnerships that made this work possible, and for their thoughtful review and comments on the manuscript. We also wish to acknowledge B. Pickering for helping to arrange antibody testing, and N. P. L. Toledo for performing the antibody testing. We are grateful for the work of D. Bulir in developing the PCR testing assay used in this study. We wish to thank L. Lazure and the staff of the Granby Zoo, the technicians of the Québec rabies surveillance program, and V. Casaubon and J. Viau of the CWHC Québec for their help with sample collection. We are also grateful for the sample collection assistance of N. Pulham, S. Konieczka, J. Adams, G. McCoy, T. McGee, L. Pollock, and K. Bennett at the Wildlife Research and Monitoring Section of the NDMNRF and L. Dougherty, L. Shirose, and M. Alexandrou at the CWHC Ontario. Finally, we wish to thank the licensed fur harvesters who submitted mink for testing.

## Funding

This work was supported by the Public Health Agency of Canada, with in-kind contributions provided by all collaborating partners.

## Notes

### Competing Interest Statement

The authors have declared no competing interest.

